# Purification of immunostimulatory glucose homopolymer from broccoli (*Brassica oleracea* var. italica)

**DOI:** 10.1101/2021.05.26.445781

**Authors:** Atsushi Miyashita, Keiko Kataoka, Toshio Tsuchida, Akihiko Ano Ogasawara, Hiroshi Hamamoto, Megumu Takahashi, Kazuhisa Sekimizu

## Abstract

We prepared broccoli (*Brassica oleracea* var. italica) neutral polysaccharides (flow-through fractions of anion exchange column chromatography from hot water extracts) from different broccoli cultivars and compared their immunostimulatory effects in the silkworm muscle contraction assay. The cultivars showed a wide range of activity, with the cultivar ‘Winter dome’ showing the highest specific activity (more than 100 times higher than curdlan). Furthermore, the active substance was purified by gel filtration column chromatography. The active substance showed heterogeneous molecular weights of more than 270 kDa. Sugar composition analysis of the purified fraction revealed that more than 95% of its sugar component was glucose, suggesting that the immunostimulatory neutral polysaccharide from broccoli cultivar ‘Winter dome’ was a homopolymer of glucose. The purified fraction also induced TNFα production in cultured mouse macrophage cells. These results suggest that the glucose homopolymer in broccoli has an immunostimulatory effect on both arthropod and mammalian immune system.

## Introduction

Innate immunity is an important biological system that plays the first line of infection defense in multicellular organisms, and its proper function is regulated by various internal and external factors. In terms of infection defense, the innate immune system can rapidly remove foreign substances such as bacteria, fungi, viruses, and toxins without the use of antibodies, and is a biological defense system that is conserved across phyla (1–5). We have previously reported that the paralytic peptide (PP) pathway, which involves muscle contraction, is one of the innate immune pathways that responds to xenobiotics in the silkworm (6,7). We also developed an experimental method to quantify the muscle contraction activity and proposed a methodology to search for natural products that stimulate innate immunity using the silkworm assay system (8,9). Using the assay system, we have found so far are β-glucan (9), acidic polysaccharide from broccoli(10), and neutral polysaccharide from broccoli (11). In our recent study on the neutral polysaccharide in the broccoli, we have found that the crude sample of broccoli neutral polysaccharides induced muscle contraction (i.e., PP activation) in the silkworm, suggesting the presence of immunostimulatory neutral polysaccharides in the broccoli (11). The effect of these polysaccharides from broccoli on innate immunity may be similar to those found in other biologically derived polysaccharides (12–15).

Of the broccoli polysaccharides with immunostimulatory activity, acidic polysaccharides have been characterized to be pectin-like polysaccharides that precipitate in ethanol and bind to diethylamino-ethyl (DEAE) cellulose column (10). However, for neutral polysaccharides that do not bind to DEAE cellulose columns, there is virtually no information on their chemical structures. Moreover, previous studies have suggested that the innate immune activating ability of hot water extracts may be different across broccoli cultivars. In the present study, we started by comparing the immunostimulatory activity of crude neutral polysaccharide fractions using 15 cultivars, where the cultivar ‘Winter dome’ showed the highest activity. For ‘Winter dome’, we further purified the active substance and characterized for their chemical structure and immunostimulatory effects on mammalian immune cells.

## Results

### Immunostimulatory effects of neutral polysaccharides from different broccoli cultivars

The immunostimulatory activities of neutral polysaccharide fraction (i.e., DEAE flow-through fraction) were determined for 15 broccoli and 2 cauliflower cultivar (Table 1). The immunostimulatory activity per sugar weight (i.e., specific activity) distributed between 260 and 11000 Units/mg, with the cultivar ‘Winter dome’ showing the highest specific activity (Table 1).

**Table 1.**
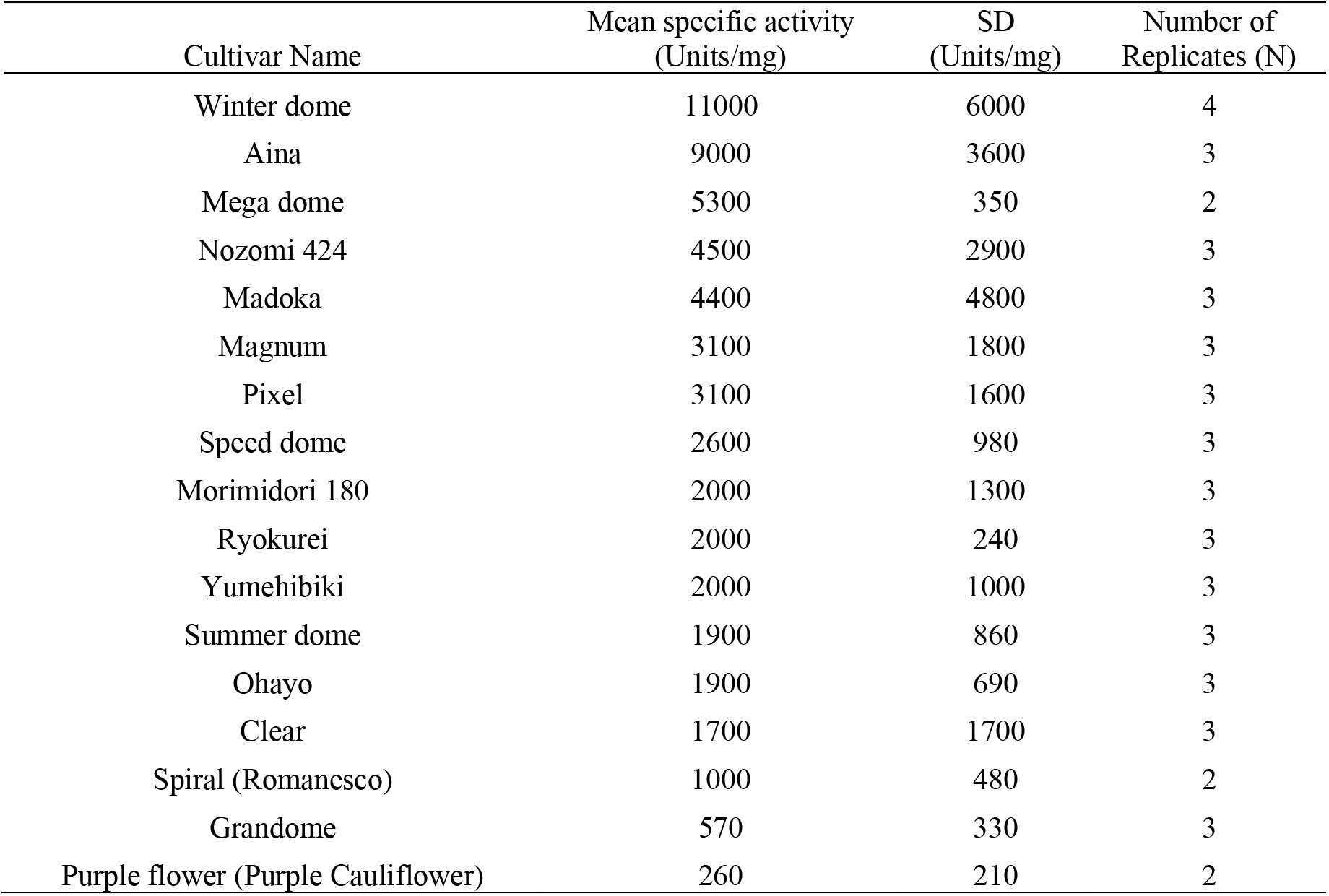
List of broccoli cultivars (including two cauliflowers).

| Cultivar Name | Mean specific activity<br>(Units/mg) | SD<br>(Units/mg) | Number of<br>Replicates (N) |
| --- | --- | --- | --- |
| Winter dome | 11000 | 6000 | 4 |
| Aina | 9000 | 3600 | 3 |
| Mega dome | 5300 | 350 | 2 |
| Nozomi 424 | 4500 | 2900 | 3 |
| Madoka | 4400 | 4800 | 3 |
| Magnum | 3100 | 1800 | 3 |
| Pixel | 3100 | 1600 | 3 |
| Speed dome | 2600 | 980 | 3 |
| Morimidori 180 | 2000 | 1300 | 3 |
| Ryokurei | 2000 | 240 | 3 |
| Yumehibiki | 2000 | 1000 | 3 |
| Summer dome | 1900 | 860 | 3 |
| Ohayo | 1900 | 690 | 3 |
| Clear | 1700 | 1700 | 3 |
| Spiral (Romanesco) | 1000 | 480 | 2 |
| Grandome | 570 | 330 | 3 |
| Purple flower (Purple Cauliflower) | 260 | 210 | 2 |

### Purification of immunostimulatory neutral polysaccharides in broccoli

Using the cultivar ‘Winter dome’, we purified the immunostimulatory substance. By the gel filtration column chromatography (we pooled Fraction II as shown in Figure 1), the specific activity increased by 3.6-fold compared to the DEAE flow-through fraction. The yield of activity was 50% (Fig. 1 and Table 2). The purification table is shown in Table 2. As shown in Figure 1, the activity appeared on different positions of the gel filtration, indicating that polysaccharides with different molecular sizes greater than 270 kDa shows the immunostimulatory effect.

**Figure 1.**
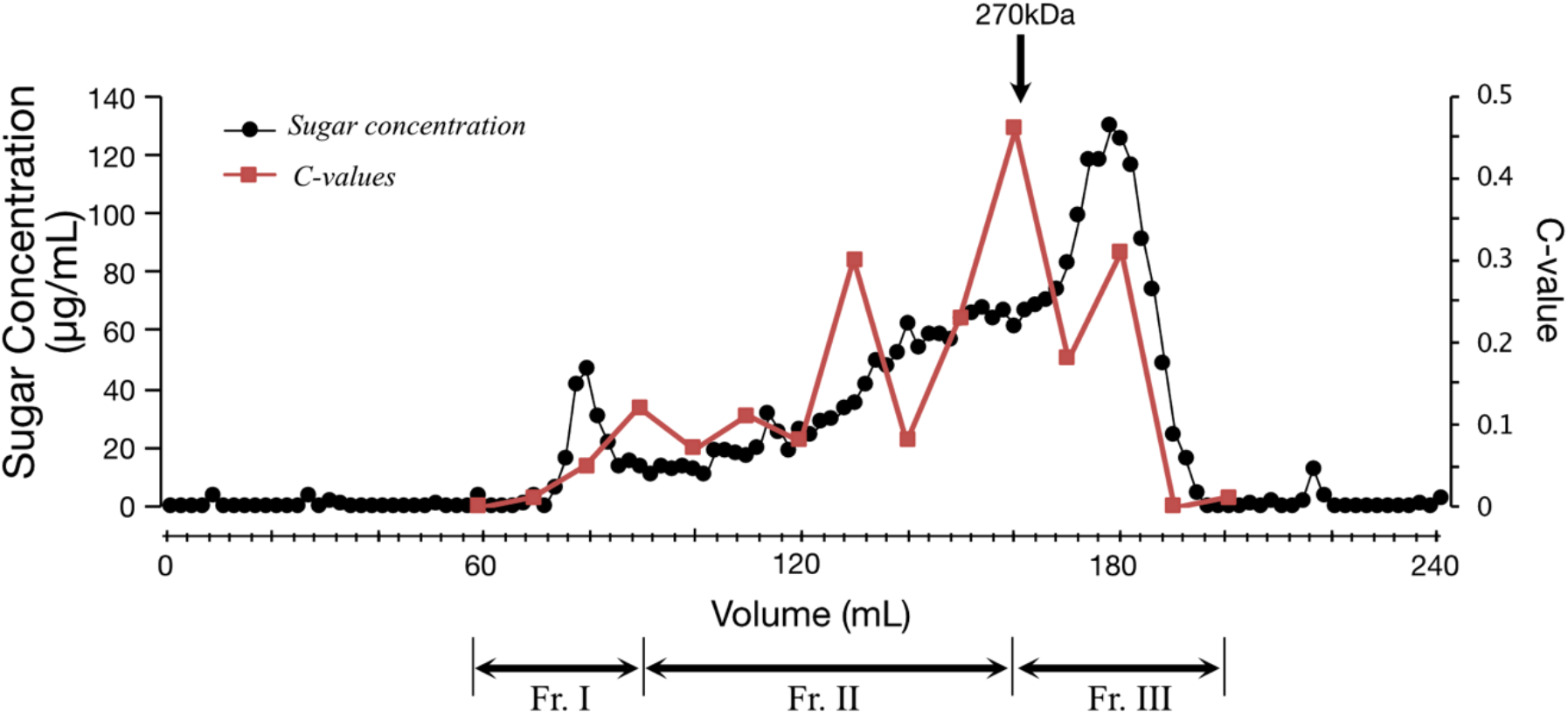
Elution profile of Sephacryl S-1000 gel filtration column chromatography. The elution profile of the gel filtration column chromatography is shown in the figure. The x-axis represents elution volume (mL), and the y-axes represent sugar concentration (μg/mL) of each fraction (the left axis) and the contraction value (the right axis). Method details for the chromatography, sugar quantification, and contraction value estimation are described in the method section. The black circles represent the sugar concentrations, and the red squares represent the contraction values. As shown in the chart, we pooled fractions into three fractions: Fr. I, II, and III. In this study, Fr. II was used for further analysis. Please also see Table 2 for the purification table.

**Table 2.** Purification Table.

| Step | Sugar Amount<br>(mg) | Total Activity<br>(Units) | Yield (%) | Specific Activity<br>(Units/mg) | Purification<br>Level |
| --- | --- | --- | --- | --- | --- |
| DEAE-FT | 5.5 | 50000 | 100 | 9000 | 1.0 |
| Gel Filtration<br>(Fr. II) | 0.77 | 25000 | 50 | 32000 | 3.6 |

### Sugar composition analysis of purified neutral polysaccharides (Fr. II)

After acid hydrolysis and ABEE labeling of Fr. II (see Fig. 1 and Table 2), we performed an HPLC analysis, where it showed a single peak derived from ABEE-labeled glucose. Other monosaccharides were below the detection limit (Fig. 2). This result indicates that the chemical structure of the neutral polysaccharide present in Fr. II is glucose homopolymers.

**Figure 2.**
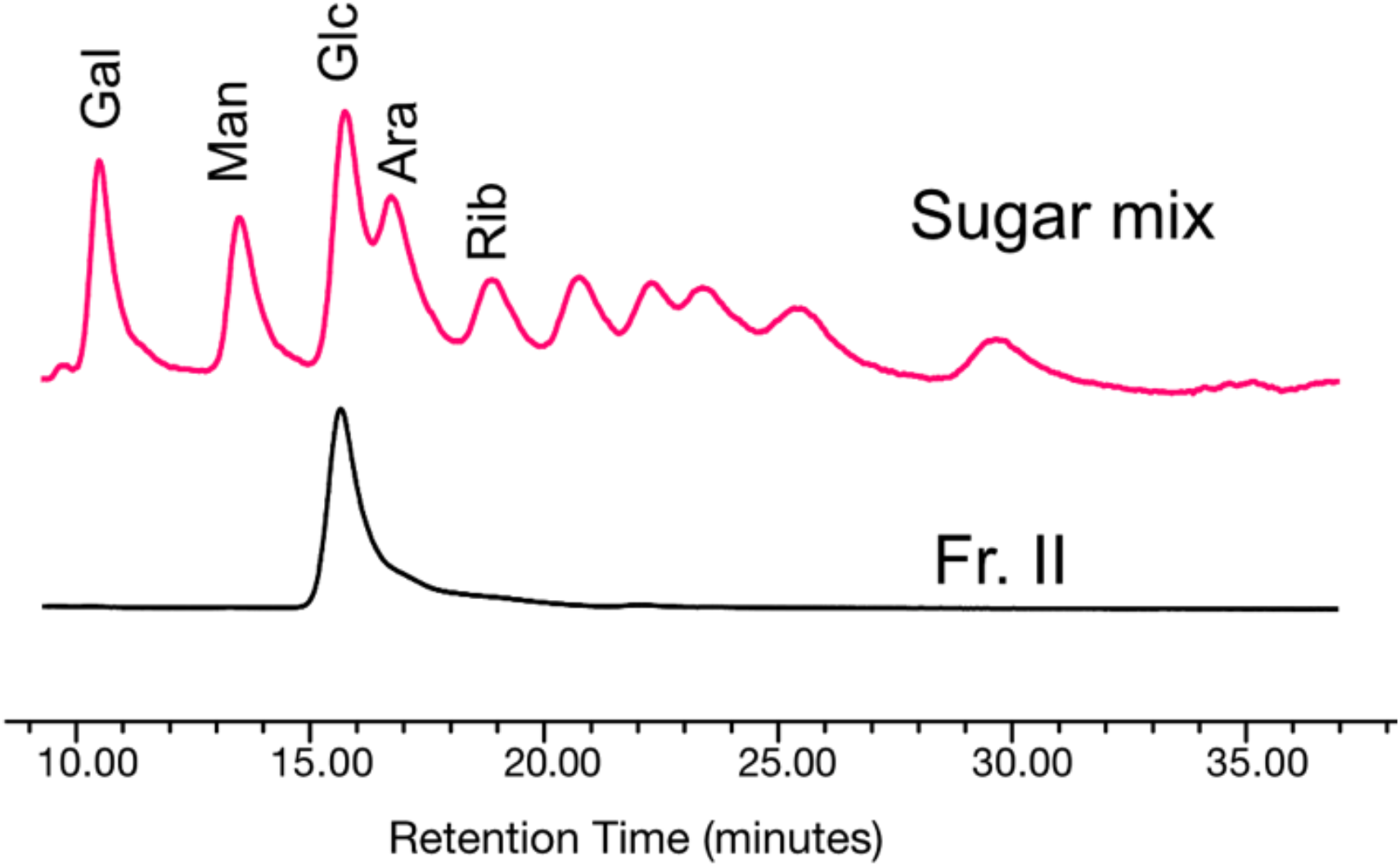
Sugar component analysis of the purified immunostimulatory neutral polysaccharide (Fr. II). Shown in the figure is the HPLC chart of sugar component analysis. The x-axis represents retention time (minutes). The red chart represents the HPLC pattern from ABEE-labelled sugar (monosaccharide) mixture. Each peak is annotated as described in the method section. The black chart represents the HPLC pattern from ABEE-labelled hydrolyzed sample (Fr. II). As shown in the figure, Fr. II showed a signal from ABEE-labelled glucose, and other monosaccharides were under the detection limit. See the method section for technical details for this experiment.

### Immunostimulatory effect of the broccoli neutral polysaccharides (Fraction II) on mammalian cells

We further assessed the immunostimulatory effect of the broccoli neutral polysaccharides (Fraction II) on mammalian cells. As shown in the Figure 3, TNF-alpha concentration in the culture medium of mouse macrophage cells was higher when treated by Fraction II (mean = 205 pg/mL, sd = 30 pg/mL) than when treated by the buffer used for gel filtration (under detection limit for all three replicates). The difference was statistically significant (p = 0.0035, Student’s t-test).

**Figure 3.**
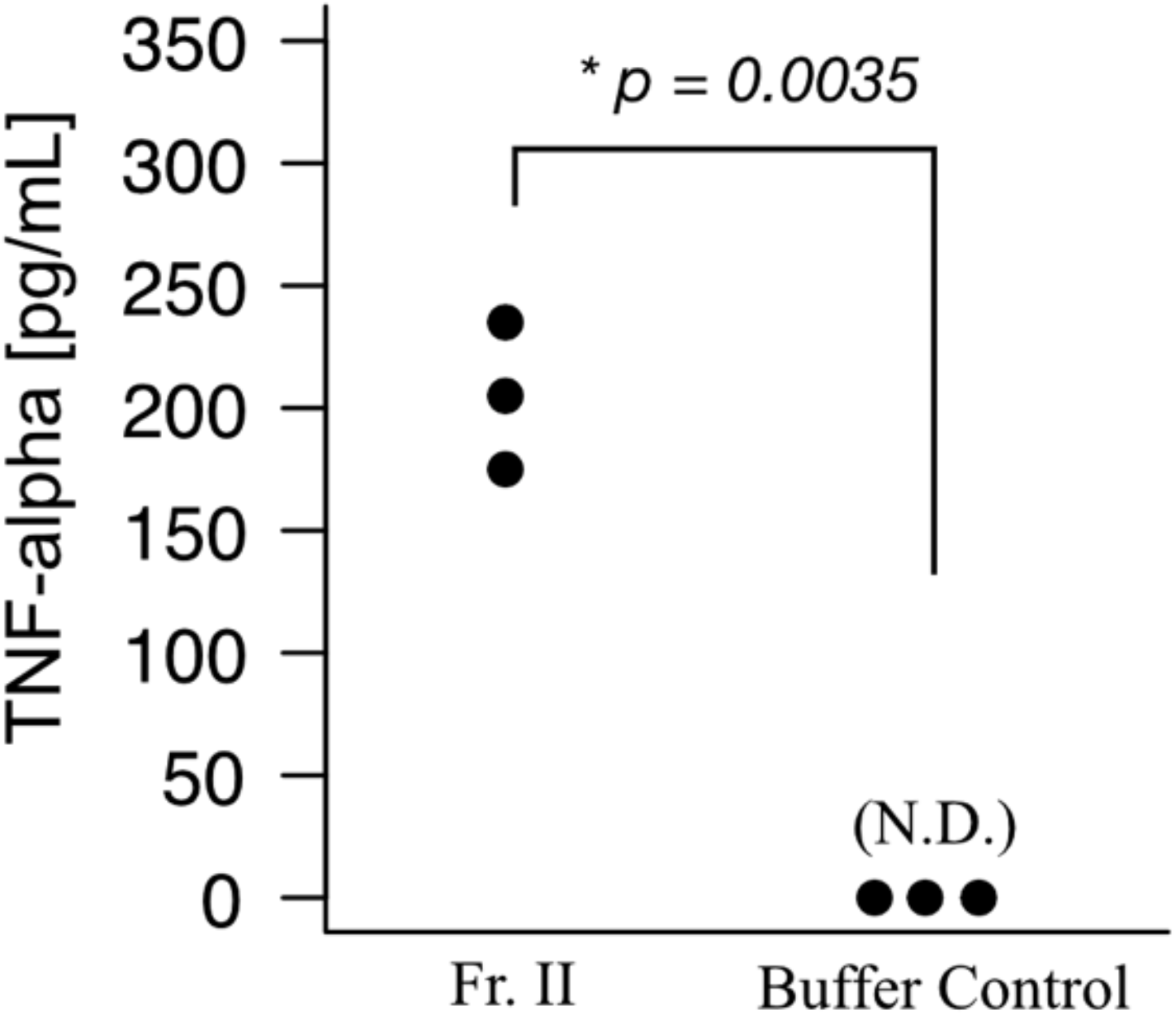
Immunostimulatory effect of the purified fraction (Fr. II) on mouse macrophage cells. Mouse macrophage cells (RAW264.7) were exposed to the sample and measured for TNF-α amount in the culture medium. The y-axis represents the TNF-α concentration (pg/mL). Same experiment was performed using the buffer that we used for the gel filtration column chromatography when we obtained Fr. II (Buffer Control), where neither of the three replicates induced TNF-α production by RAW.264.7 cells. N.D., not detected. There was a statistically significant difference between the two groups (p = 0.0035, Student’s t-test).

## Discussion

In this study, we have shown that neutral polysaccharides (glucose homopolymers) present in the edible portion of broccoli can activate the innate immune system. This activity was verified in this study by both the silkworm muscle contraction assay and the TNF-α induction assay using mammalian macrophage cells. The immunostimulatory effects of neutral polysaccharides of broccoli were reported in our previous report (11), and in this study, we have further purified the active substance and obtained the information about its chemical structure based on the molecular size estimation in gel filtration column chromatography and the results of sugar composition analysis. In plants and fungi other than broccoli, polysaccharides from edible materials such as *Cordyceps militaris* (12), *Mosla chinensis* (13), *Panax ginseng* (14), and *Dimocarpus longan* (15) have been reported to have immunostimulatory effects. Nevertheless, information about their chemical structure is scarce.

In the present study, we have also compared the immunostimulatory activity (i.e., silkworm muscle contraction) of the DEAE flow-through fractions from 17 broccoli and cauliflower cultivars. The results demonstrated differences among the cultivars for the specific activity (i.e., amount of activity per sugar weight). The difference in immunostimulatory activity among the cultivars was pointed in the previous report (11), and whether this difference is due to the amount of active substance or to the presence of active substances with different chemical structures needs further investigation. The DEAE flow-through fraction of the cultivar ‘Winter dome’, which showed the highest specific activity (Table 1), was further purified by gel filtration. As a result, we found that the active substance in the purified fraction (Fr. II) showed heterogeneous molecular weights larger than 270 kDa and a specific activity of 32,000 Units/mg. We have been using silkworm muscle contraction as an indicator to identify innate immune activators in various food materials, and this value (i.e., 32,000 Units/mg) seems outstanding. For example, the specific activity of the neutral polysaccharide fraction of broccoli is more than 300 times higher than that of curdlan, which has a β-(1,3)-glucan (glucose homopolymer) structure and has been regarded as one of the most active polysaccharides for immunostimulatory effect. There may be a unique chemical structure of glucose homopolymer that shows such outstanding specific activity.

Two groups of eukaryotes, fungi and plants, possess large amounts of polysaccharides in a structure called the cell wall. Polysaccharide recognition in eukaryotes plays an important role in defense against pathogenic fungi. Lectins and complement proteins are known as molecules that recognize polysaccharides in fungi (19). It has been reported that α-(1,3)-glucan produced by *Histoplasma capsulatum* contributes to virulence by hiding immunostimulatory β-glucans from detection by host phagocytes (20). In fact, in our assay system for measuring immune activation ability using silkworm muscle contraction as an indicator, we reported that the strength of activity varied depending on the origin and chemical structure of β-glucans (9). Further investigation will open up our understanding of the interaction between diverse polysaccharide substances and the animal immune system.

## Experimental procedures

### Broccoli

Fifteen cultivars of broccoli (*B. oleracea* var. italica) and two cultivars of cauliflower (*B. oleracea* var. botrytis) were provided by the National Agriculture and Food Research Organization (Tsukuba, Japan) (16). Spring crops were used in this study. The list of cultivars is shown in Table 1.

### Hot water extract of broccoli

Hot water extracts of broccoli were prepared by a method previously reported (11). Briefly, flower buds were cut with scissors, boiled in water for 2 minutes, and then immersed in ice-cold water to cool well. Fifteen grams (wet weight) of buds were ground in 30 mL of pure water using a polytron. The sample was then autoclaved at 121°C for 20 minutes and was spun at 8,000 rpm for 10 min to obtain the supernatant.

### Silkworm muscle contraction assay

Decapitated silkworm specimens were prepared using rearing 5-day-old (days were counted from the final molt) 5^th^ instar silkworms (17). Silkworms were fed artificial diet containing antibiotics (17). Fifty microliter of the sample solution was injected into the silkworm using tuberculin syringe with a 27-gauge needle, and the body length was monitored over 10 minutes after the sample injection. The contraction value C=(x-y)/x was calculated for the length x cm of the silkworm at the beginning of the experiment and the length y cm at the maximum contraction. The contraction values for different sample doses were plotted, and the sample volume with a contraction value C of 0.15 was determined, and its activity was set to 1 unit (17).

### Purification of immunostimulatory substance from broccoli

#### I. DEAE Cellulose column chromatography

Two volumes of PCI (phenol/chloroform/isoamyl alcohol) was added to the hot water extract of broccoli, and the aqueous layer was collected after centrifugation (8,000 rpm, 5 minutes). Two volumes of ethanol was added to the aqueous layer, and the precipitate formed by centrifugation (8,000 rpm, 10 min) was dissolved in 10 mM Tris-HCl (pH 7.9) buffer, and loaded to a DEAE cellulose column (GE Healthcare, bed volume = 20 mL) equilibrated by the same buffer, and we collected the flow-through fraction.

#### II. Gel filtration column chromatography

The flow-through fraction from DEAE column chromatography was concentrated by ethanol precipitation and then applied to a gel filtration column (Sephacryl S-1000, diameter = 15 mm, height = 1080 mm, bed volume = 190 mL). The column was equilibrated by 10 mM Tris/HCl (pH=7.9) buffer in advance and the same buffer was used for elution. The fraction size was 2 mL/fraction, and each fraction was subjected to sugar quantification and silkworm muscle contraction assay. Sugar quantification was performed according to a previously reported method (17). Active fractions were pooled to obtain three fractions (fractions I, II, and III). We used fraction II for further characterization, as the fraction showed an increased specific activity (please also see the purification table as shown in Table 2).

### Sugar Composition Analysis

Sugar composition analysis was performed using 4-aminobenzoic acid ethyl ester (ABEE) labeling method as previously reported (18). After acid hydrolysis of the purified polysaccharide fraction (fraction II), the ABEE-labeled samples were analyzed by an HPLC using a reversed-phase column. Monosaccharide standards were used as control, and each monosaccharide was identified based on elution time. To further validate the peak annotation, the samples were mixed with ABEE labeled glucose samples and analyzed by HPLC, where we confirmed that the peaks overlapped for each sample.

### TNF-alpha induction assay using murine macrophage cells

Experiments were done as described in the previous work (18). In a 96-well plate, we spread RAW264.7 cells (1×10^5^ cells/well) in RPMI 1640 (+10% FBS) and cultured for 24 hours at 37°C. After removing the culture medium, we immediately put the purified sample (Fraction II) or the buffer used for the chromatography (10mM Tris-HCl, pH = 7.9) suspended in RPMI 1640 (non-FBS) medium. We then incubated the cells for 6 hours. The final sugar concentration added in the medium for Fraction II-treated group was 22 μg/mL. We collected the cell culture supernatant and measured the TNF-α concentration using Mouse TNF-α Quantikine ELISA Kit MTA00B (R&D SYSTEMS).

## Data availability

All data are contained within the manuscript

## Acknowledgement

This work was supported by a JSPS KAKENHI (Grant# 20K16253) to AM, and a JSPS KAKENHI (Grant# 21H02733) to KS. We thank Genome Pharmaceuticals Institute, Co. Ltd., for technical supports.

## Author contributions

Conceptualization: KS

Data Curation: AM, KK, TT, AAO

Formal Analysis: AM

Funding acquisition: AM, KS

Resources: MT

Writing – original draft: AM

Writing – review & editing: AM, HH, MT, KS

## Abbreviations and nomenclature

ABEE: 4-aminobenzoic acid ethyl ester
PP: Paralytic peptide
DEAE: Diethylaminoethyl
TNF: Tumor necrosis factor
HPLC: High performance liquid chromatography

